# Salactin, a dynamically unstable actin homolog in Haloarchaea

**DOI:** 10.1101/2023.03.09.531933

**Authors:** Jenny Zheng, Alex Lammers, John Mallon, Thomas Litschel, Edmund R.R. Moody, Diego A. Ramirez-Diaz, Amy Schmid, Tom A. Williams, Alexandre W. Bisson-Filho, Ethan Garner

## Abstract

Across the domains of life, actin homologs are integral components of many essential processes such as DNA segregation, cell division, and cell shape determination. Archaea genomes, like those of bacteria and eukaryotes, also encode actin homologs, but much less is known about these proteins’ in vivo dynamics and cellular functions. We identified and characterized the function and dynamics of Salactin, an actin homolog in the hypersaline archaeon Halobacterium salinarum. Despite Salactin’s homology to bacterial MreB proteins, we find it does not function as a MreB ortholog in H. salinarum. Rather, live-cell imaging revealed that Salactin forms dynamically unstable filaments that grow and shrink out of the cell poles. Like other dynamically unstable polymers, Salactin monomers add at the growing filament end and its ATP-bound critical concentration is substantially lower than the ADP-bound form. When H. salinarum’s chromosomal copy number becomes limiting under low phosphate growth conditions, cells lacking Salactin show perturbed DNA distributions. Taken together, we propose that Salactin is part of a previously unknown chromosomal segregation apparatus required during low-ploidy conditions.

## Introduction

Actin and its homologs are present in all domains of life and are involved in various processes, such as cell motility, division, shape determination, and DNA segregation (Dunleavy et al., 2019; Eun et al., 2015; Fuesler & Li, 2012; Jékely, 2014). Eukaryotic actin is critical for the motility, division, and shape of eukaryotic cells (Dominguez & Holmes, 2011; Pollard & Cooper, 2009). Bacteria also contain actin homologs that are involved in a variety of functions: 1) MreB creates and maintains rod shape (Jones et al., 2001) 2) MamK positions magnetosomes along the cell length (Komeili et al., 2006; Scheffel et al., 2006) 3) FtsA is a central component of the division machinery (Haeusser & Margolin, 2016) and 4) plasmid-encoded actin proteins such as AlfA and ParM ensure low-copy plasmid inheritance (Derman et al., 2009; Møller-Jensen et al., 2002, 2003; Polka et al., 2009, 2014). Phylogenomics identified several actin and tubulin homologs in archaea (Makarova & Koonin, 2010; Spang et al., 2015; Yutin & Koonin, 2012; Zaremba-Niedzwiedzka et al., 2017). Two archaeal tubulin homolog families (FtsZ and CetZ) have been visualized in the haloarchaeon *Haloferax volcanii* and determined to be involved in division and cell shape, respectively (Duggin et al., 2015; Liao et al., 2021).

Comparatively, the functions of the archaeal actins have been far less studied: we still lack an understanding of the *in vivo* dynamics or function of any archaeal actin. Various studies have made progress in our understanding: Crenactin’s (from Crenarchaeota) localization and correlation with cell shape suggests it could be involved in cell shape formation (Ettema et al., 2011; Izoré et al., 2016; Lindås et al., 2014). Bioinformatics identified multiple actins in the TACK and Asgard families with high sequence similarity to eukaryotic actin (Cai et al., 2020; Imachi et al., 2020; Liu et al., 2021; Seitz et al., 2019; Spang et al., 2015; Zaremba-Niedzwiedzka et al., 2017; Zhang et al., 2021), and the Asgard genomes also encode multiple eukaryotic-like actin-modulating proteins (Akıl et al., 2020, 2022; Akıl & Robinson, 2018; Eme et al., 2017; Stairs & Ettema, 2020; Survery et al., 2021). Accordingly, cryo-electron microscopy of Loki Asgard archaea revealed long eukaryotic-like actin filaments enriched within cellular protrusions (Rodrigues-Oliveira et al., 2022).

Motivated by previous bioinformatic analyses suggesting haloarchaea contained bacterial MreB-like actin homologs (Makarova et al., 2010), we searched for actins in the archaeal model *Halobacterium salinarum*. Here, we identified and characterized a MreB homolog in *H. salinarum* which we name Salactin.

## Results

### Identification of Salactin

To search for putative actin fold proteins suggested by Makarova and colleagues, we used the Escherichia coli’s MreB protein sequence as input to JackHMMER, a statistical tool based on Hidden Markov Models (HMM) (Johnson et al., 2010). Limiting the results to archaeal proteins, we initially obtained 273 candidates across multiple phyla, but most were annotated as DnaK or other chaperones. However, a second JackHHMER iteration revealed an additional 86 candidates whose majority (58 out of 86) grouped within the Halobacteria class. In *H. salinarum*, we identified a conserved putative MreB (GenBank AAG18772.1), which we named Salactin.

Phylogenetic analysis of *salactin* (Fig. 1A, Fig. S1) indicated that the gene is broadly conserved across the Haloarchaea, with closely related homologs also present in the other lineages of Methanotecta (the euryarchaeotal clade comprising Haloarchaea, Methanomicrobia, Methanocellales, Methanophagales, and Archaeoglobi) (Adam et al., 2017). Phylogenic analysis (Fig. 1A) suggests that *salactin* was already present at the root of Methanotecta. However, while Salactin is more closely related to MreB than to any other actin subfamily, a high level of sequence divergence from other MreB homologs makes it difficult to place Salactin within the broader diversity of MreB sequences. The most closely related sequences outside Methanotecta are divergent MreB homologs in some other Euryarchaeota (Poseidonales) and Bacteria (Fig. S1), making it difficult to determine the history of Salactin prior to the common ancestor of Methanotecta.

**Figure 1:**
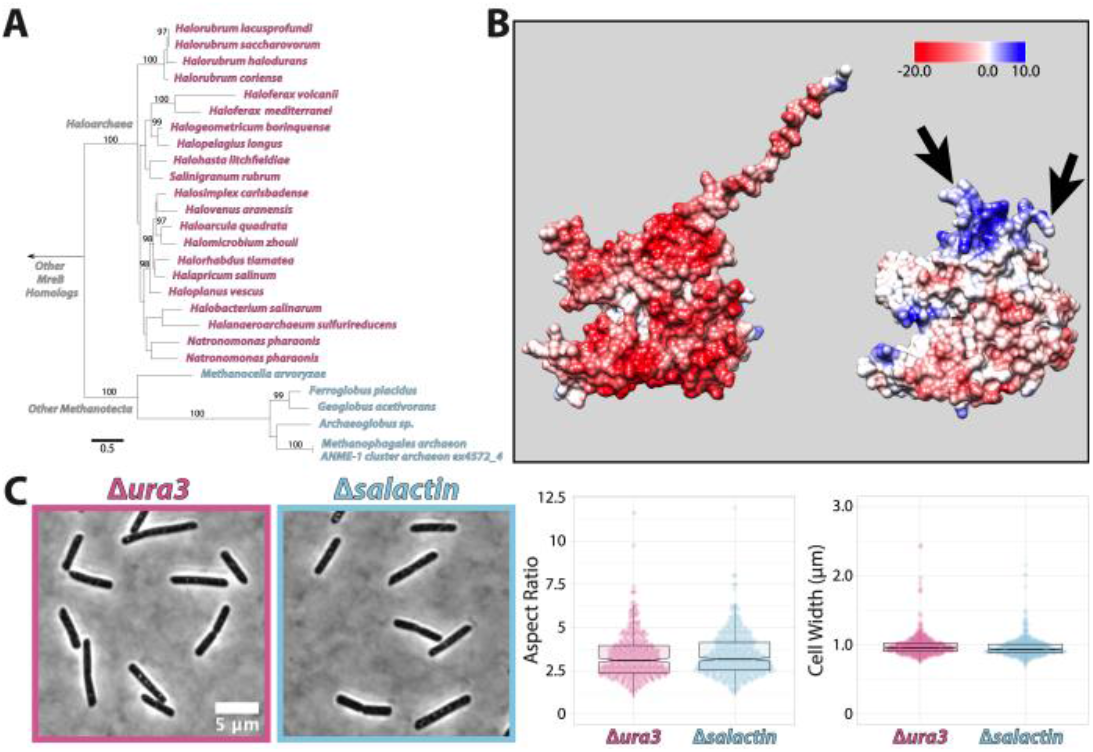
(A) Tree of the representative species in the Methanotecta clade in the Euryarchaeota phylum with a closely related homolog of Salactin. The numbers on branches are the ultrafast bootstrap support and represent how reliable the branch is, where 100% is well supported, and greater than 95% is strongly supported. (B) Alphafold2 predicted structures of Salactin (left) and MreB from Bacillus subtilis (right) colored by their surface electrostatic potential. Arrows indicate regions of MreB that have been implicated in membrane binding. (C) Phase images of Δura3 parent strain (pink) and Δsalactin (blue) H. salinarum cells in rich media (CM+URA) (left) showing that both exhibit the same rod shape. Violin plot of the aspect ratio (mid) and width (right) of Δsalactin cells and Δura3 cells demonstrates no statistically significant difference between strains (width p = 0.0016, aspect ratio p = 0.22).

### Salactin does not function as a MreB ortholog

Salactin’s sequence contains the characteristic nucleotide binding motifs found within HSP70/actin fold proteins (Bork et al., 1992). Structural predictions by AlphaFold2 (Jumper et al., 2021) suggested a canonical actin fold (AlphaFold Database Q9HSN1) with a long, highly charged disordered tail on the N-terminus (Fig. 1B). While Salactin’s predicted structure resembles MreB’s, it lacked any of the amphipathic helices or surface-exposed hydrophobic residues that are required for MreB’s membrane binding (Mao et al., 2022; Salje et al., 2011), suggesting Salactin does not directly associate with the membrane.

MreB is involved in establishing and maintaining the rod shape of many bacteria (Jones et al., 2001; Shi et al., 2018; Yulo & Hendrickson, 2019). To determine if Salactin acts as a MreB ortholog and is required for *H. salinarum’s* rod shape, we created a strain where *salactin* was deleted in a Δ*ura3* background (Δ*salactin*Δ*ura3* referred to as Δ*salactin)* and a Δ*ura3* strain that we used throughout this work as our control. Because *H. salinarum* is highly polyploid, we performed whole-genome sequencing and determined that Δ*salactin* cells are devoid of *salactin* sequences, and therefore deleted from all chromosomal copies. Furthermore, we confirmed that no second site suppressor mutations were detected in comparison to the Δ*ura3* sequences, thereby ruling out *salactin* essentiality in rich media (Table S1).

Single-cell image analysis from phase-contrast microscopy revealed that Δ*salactin* cells in exponential phase have rod shapes that are indistinguishable from Δ*ura3* cells, with no significant difference in width, aspect ratio, area, length, or circularity relative to Δ*ura3* cells (all p-values > 0.0016: width p = 0.0016, aspect ratio p = 0.22, area p = 0.21, length p = 0.8774, circularity p = 0.34) (Fig. 1C, Fig. S2). As deletion of *salactin* has no effect on the rod shape of *H. salinarum*, these experiments suggested that Salactin does not functionally act as an MreB ortholog.

### Salactin filaments display dynamic instability *in vivo*

To gain further insight into Salactin’s function, we examined its localization and dynamics *in vivo*. We created constructs by fusing Salactin to either monomeric superfolder GFP (msfGFP) or HaloTag and driven by a constitutive, strong ribosomal promoter (Prpa) on a high-copy plasmid. By expressing Salactin-msfGFP ectopically (strain (hsJZ52) from a plasmid along with the native copy of *salactin*, we visualized Salactin-msfGFP’s localization and dynamics using time-lapse fluorescence microscopy. These experiments did not reveal short filaments of Salactin moving around the cell width as one would expect for an MreB homolog (Domínguez-Escobar et al., 2011; Garner et al., 2011; van Teeffelen et al., 2011). Instead, Salactin filaments showed a very different phenotype: in 74.55% of cells, Salactin polymers grew out of the cell poles toward midcell, then would suddenly depolymerize, with filaments rapidly shrinking back to the poles (Fig. 2A, SM1, Table S2). The stochastic switching between steady elongation and rapid depolymerization is known as dynamic instability (Mitchison & Kirschner, 1984). Dynamic instability can be seen in kymographs created with a line from the pole to the mid-cell, where dynamic instability creates right triangles in the kymograph (Fig. 2B): the hypotenuse is the slow phase of polymerization, and the adjacent side arises from rapid depolymerization (Zwetsloot et al., 2018). We observed that this behavior of Salactin would occur multiple times from one pole during a 1-hour observation, shown by the repeating triangles in the kymographs. Analysis of 50 kymographs indicated Salactin filaments grow at a rate of 3.88nm/s (Fig. 2C), which (assuming a double-stranded filament and a dimer subunit rise of 5nm) would cause the filament to grow by 1.5 monomers each second. In contrast, all filament catastrophes occurred within one frame, a rate so fast we could not measure the actual depolymerization rate. By examining the longest filament that depolymerized within a single frame we could estimate Salactin’s minimal depolymerization rate to be 600nm/s or 240 monomers per second. The distribution of filament lengths and time before catastrophe had a mean of 2.02µm and 449.2 seconds, respectively (Fig. 2C). The distribution of the time before catastrophe appears to have an exponential tail (R-squared = 0.9643 and 0.9335 for exponential fit and linear fit to log frequency, respectively) (Fig. S3), in line with the expected distribution of a broad class of models of dynamic instability (Dieterle et al., 2022).

**Figure 2:**
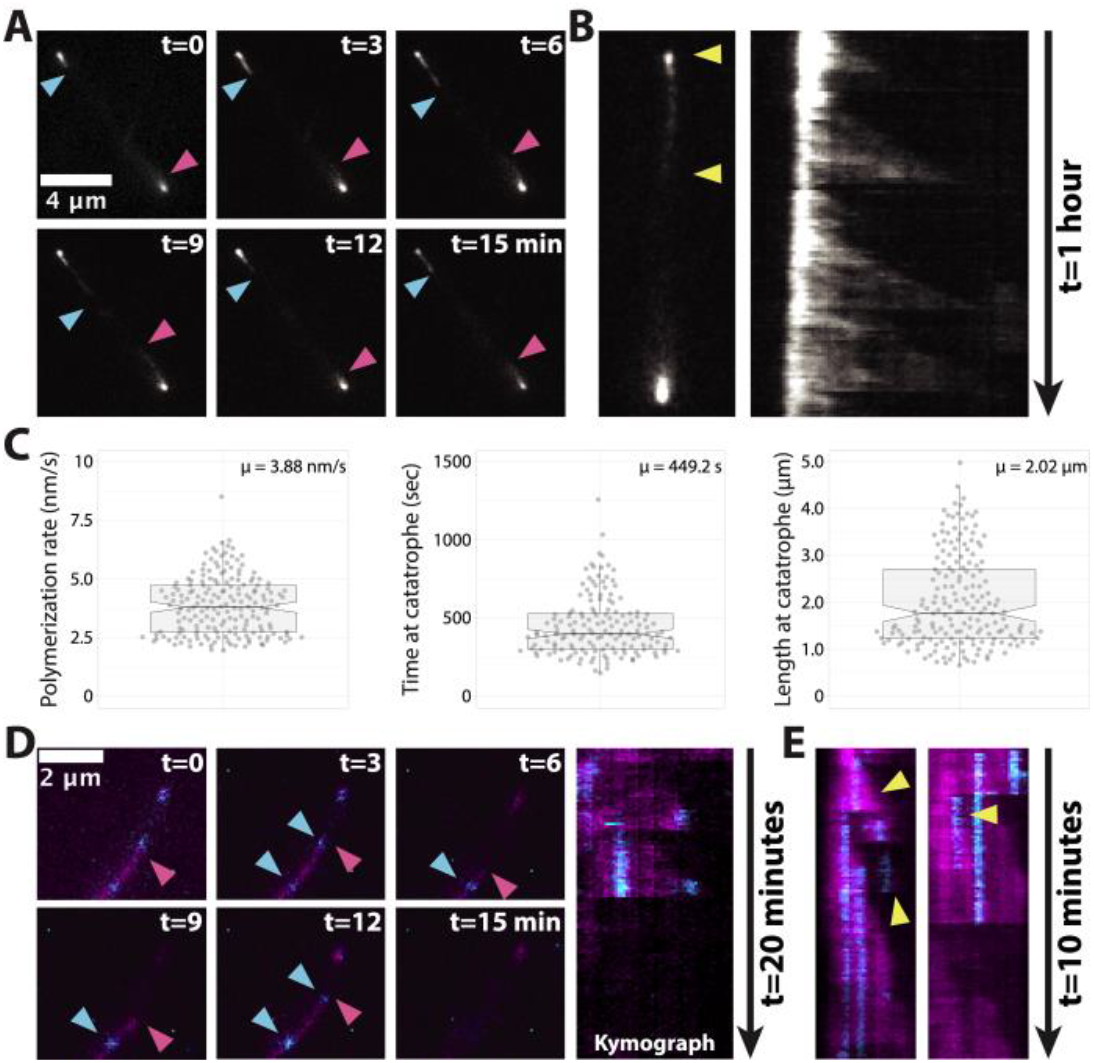
Characterization of in vivo Salactin dynamics. (A) A montage of Salactin-msfGFP in a H. salinarum cell (strain hsJZ52). The two arrowheads indicate each end of the filament. (B) A representative cell (left) used to create a kymograph (right). Yellow arrowheads indicate the region used for drawing the corresponding kymograph. (C) Violin plot of measured in vivo polymerization rates (left, N = 184), time until catastrophe (mid, N = 152), and length at catastrophe (right, N = 181) obtained from analysis of 50 kymographs. (D) A montage of a speckle labeled Salactin-HaloTag filament in a H. salinarum cell (strain hsJZ86) (left). The entire Salactin-HaloTag polymer was labeled with JF505 (magenta), and also sparsely labeled with JF549 to generate speckles (cyan). Red arrowhead indicates the filament end, blue arrowhead indicates a single molecule. Kymograph for the filament trace showing that monomers remain stationary within the growing filament (right). (E) Two example kymographs showing multiple triangles (indicated by yellow arrowheads) arising from diffraction-limited filaments, indicating these structures may be composed of multiple filaments.

We note these measurements may not be exactly the same as untagged Salactin filaments inside the cell, as (similar to many other biological polymers) fluorescent fusions to Salactin appear to affect its polymerization: when Salactin was fused to HaloTag and expressed ectopically from a plasmid in the presence of the native copy of *salactin* (strain hsJZ86), dynamically unstable filaments were observed in 61.18% of cells (Fig. 2D, Table S2, SM2, SM3, Fig. S4A). However, cells expressing Salactin-HaloTag as the sole copy (strain hsJZ106) showed a considerable reduced fraction of cells showing dynamic instability (38.76%) (Table S2, SM4, Fig. S4B). Likewise, cells expressing Salactin-msfGFP as the sole copy (strain hsJZ95) in the cell showed filaments in only 2.4% of cells, and none of these filaments showed any dynamics (Table S3, SM5, Fig. S4C). Also, Salactin-HaloTag fusions expressed as the only copy in the Δ*salactin* background (strain hsJZ106) yielded a different polymerization rate than when Salactin-HaloTag was expressed ectopically in wild-type cells in addition to the native copy (strain hsJZ86) (Fig. S4D).

### Salactin monomers are added at the growing filament end

To determine where Salactin monomers are added into growing filaments, we “speckle labeled” Salactin-HaloTag filaments by incubating cells with two Janelia Fluor dyes, JF549 at very low levels (to speckle filaments) and JF505 at much higher levels (to label the rest of the filament). Timelapse microscopy of these cells revealed that the JF549 speckles were stationary within growing Salactin filaments: speckles appeared as filaments grew and remained in the same place until they disappeared when the filament depolymerized (Fig. 2D, SM2). This demonstrates that new monomers are added to the end of the growing filament rather than being added at the filament ends at the cell poles. In addition, kymographs showed multiple depolymerization events within what appeared to be a single filament, suggesting that some of the diffraction limited polymers may contain multiple filaments (Fig. 2E), similar to what was observed with ParM filaments in *Escherichia coli* (Campbell & Mullins, 2007; Salje & Löwe, 2008). Taken together, this data demonstrates that Salactin forms dynamically unstable filaments inside *H. salinarum*.

### Salactin polymerization *in vitro*

To further understand Salactin, we purified Salactin to examine its polymerization *in vitro* (Fig. S5). Given *H. salinarum* is a halophile, we first optimized buffer KCl concentration using a malachite green assay to indirectly measure polymerization. This revealed Salactin’s ATPase activity increased with increasing salt (Fig. 3A, Fig. S6), an unsurprising result given the cytoplasm of *H. salinarum* contains ∼4.5 M KCl (Oren, 2006; Strahl & Greie, 2008). Given that the solubility of KCl is ∼3.55 M at room temperature (Pinho & Macedo, 2005), we used the highest KCl concentration (2.9 M) that yielded 1) the highest ATPase activity and 2) reproducible results without precipitation in our assays.

**Figure 3:**
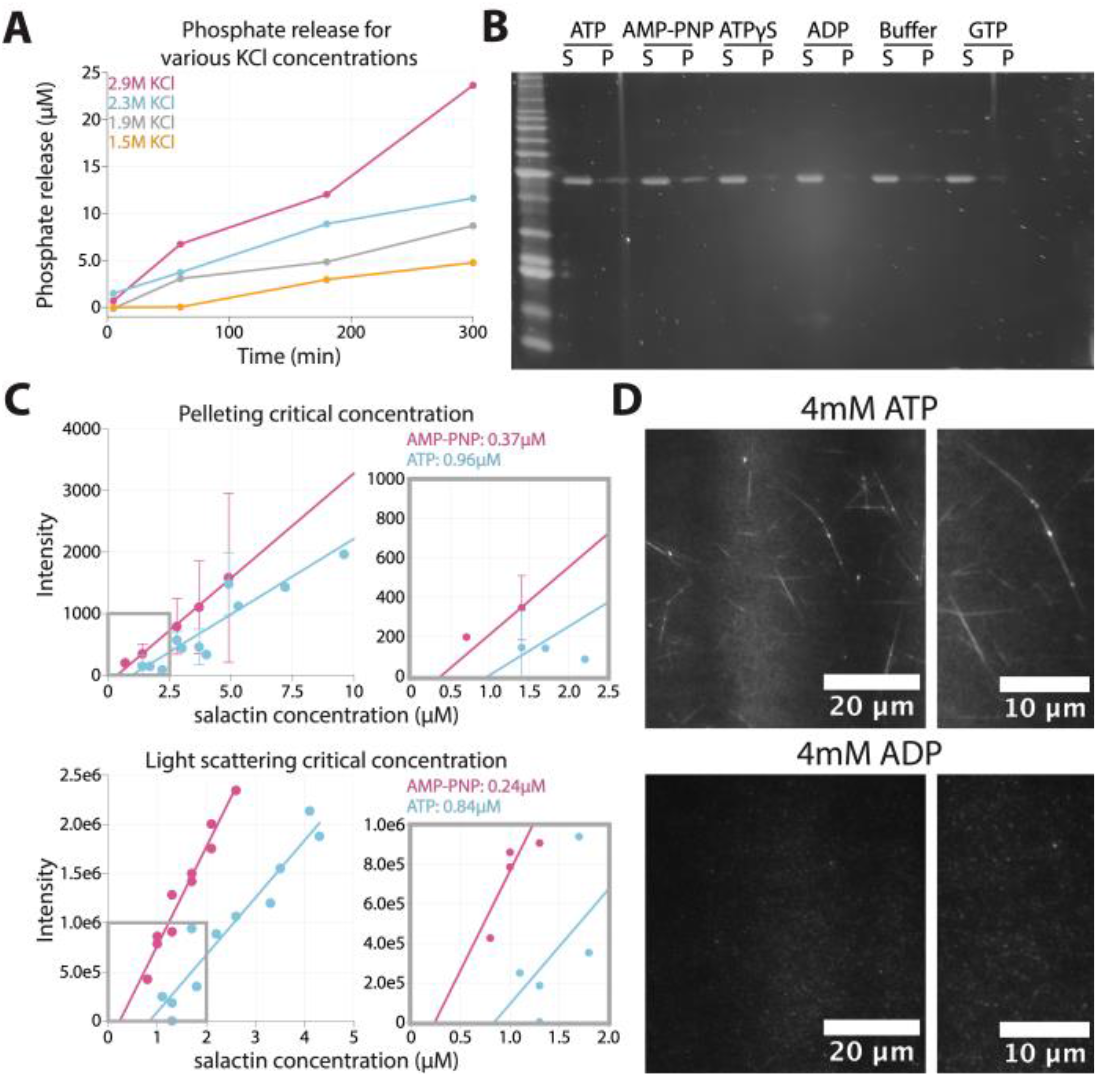
in vitro polymerization of Salactin. (A) Malachite Green Assay using 4 µM Salactin in different salt conditions (1.5M, 1.9M, 2.3M, 2.9M). The higher ATPase activity implies polymerization is favored at higher salt concentrations. (B) Salactin polymerization only occurs in the presence of ATP and AMP-PNP. Salactin polymerization was assayed by pelleting, using 2.5mM ATP, ATP analogs, ADP, or GTP. Gels were stained with SYPRO Orange. S = supernatant, P = pellet. (C) Critical concentrations of Salactin determined by pelleting (top) and light scattering (bottom). (D) Polymers of Salactin mixed with cy3B-conjugated Salactin-GSKCK are seen with TIRF microscopy in the presence of ATP but not ADP. Right panels are zoomed-in images of the left panel.

We first conducted pelleting assays to quantify Salactin’s *in vitro* polymerization. This revealed that 4.9 µM of Salactin polymerizes in the presence of ATP and AMP-PNP but not in the presence of ADP, GTP, or ATPγS (Fig. 3B). Next, we measured the critical concentrations of Salactin in the presence of different nucleotides, as dynamic instability arises from a substantial difference in the dissociation constants of the ATP and ADP-bound monomers for filament ends (Garner et al., 2004; Petek et al., 2017). We measured Salactin’s ATP and AMP-PNP critical concentrations using both pelleting and right-angle light scattering. Both assays gave similar values: pelleting yielded a critical concentration of 0.96 µM for ATP and 0.37 µM for AMP-PNP, and light-scattering yielded 0.84 µM for ATP and 0.24 µM for AMP-PNP (Fig. 3C, Fig. S7). Importantly, we were unable to observe any polymerization of Salactin in the presence of ADP up to 10 µM Salactin protein (Fig. S8), indicating that ADP bound Salactin has a much higher critical concentration than the non-hydrolyzed ATP bound state (0.37 µM). Similar to what was observed with ParM and Alp7a (Garner et al., 2004; Petek et al., 2017), the intermediate critical concentration of ∼0.9 µM in the presence of hydrolyzable ATP likely reflects the “emergent critical concentration,” the free monomer concentration that arises from the relative proportion of growing and depolymerizing filament ends. Together, these experiments indicate that, under our buffer and salt conditions, ATP and ADP-bound Salactin monomers have at least a 30-fold difference in their affinity for filament ends, giving the large energetic differential required for dynamic instability (Garner et al., 2004; Petek et al., 2017).

We attempted to visualize Salactin filaments using negative stain electron microscopy, but the high salt in our buffer caused crystalline precipitates, inhibiting the observation of filaments, a common issue that arises when high salt is used with negative staining (Scarff et al., 2018). We next attempted to visualize filaments *in vitro* using Total Internal Reflection Fluorescence (TIRF) microscopy. For this, we mixed 7 µM unlabeled Salactin with 0.34 µM cy3B-conjugated Salactin in the presence of 3 mM ATP. No filaments were observed under these initial conditions, but long Salactin bundles were observed in the presence of ATP but not ADP when we increased the macromolecular crowding with 17% PEG (Fig. 3D). These bundles did not display any depolymerization or dynamic instability during 1 hour of imaging, possibly caused by the crowding agents stabilizing filaments (Demosthene, 2021; Minton, 2000), or from the different between the intercellular KCl concentration relative to the concentration in our buffers, as past work has shown that increasing salt changes the strength of hydrophobic and ionic interactions, which can affect interactions between monomers (Kang et al., 2013).

### Salactin influences viability and DNA partitioning under low phosphate growth conditions

As our *in vivo* data demonstrates Salactin does not function as an MreB, we conducted other phenotypic tests to gain insight into Salactin’s function. First, we saw no difference in the growth rates of Δ*salactin* and Δ*ura3* parent strains growing in rich media, which we will hereafter call “standard phosphate” media (Fig. S4A). Likewise, we saw no statistical difference in the motility between these strains (Fig. S9B).

Further analysis into *salactin*’s genomic context showed poor synteny conservation even within the Halobacterium genus (Fig. 4A), making it difficult to ascribe any function from its genetic neighborhood alone. However, in *H. salinarum* and its closest relatives, the *salactin* gene is in proximity to multiple genes involved in DNA replication and repair, suggesting that Salactin could be involved in a DNA related process. As Haloarchaea are highly polyploid (Barillà, 2016; Soppa, 2022), any phenotypes arising from a DNA related process might not manifest unless ploidy is reduced. Past work has shown the ploidy of *Haloferax volcani* drastically decreases (from 30 down to 2) when cells are grown in low phosphate media, suggesting cells might limit and scavenge the excess chromosomal copies to increase their viability (Zerulla et al., 2014).

**Figure 4:**
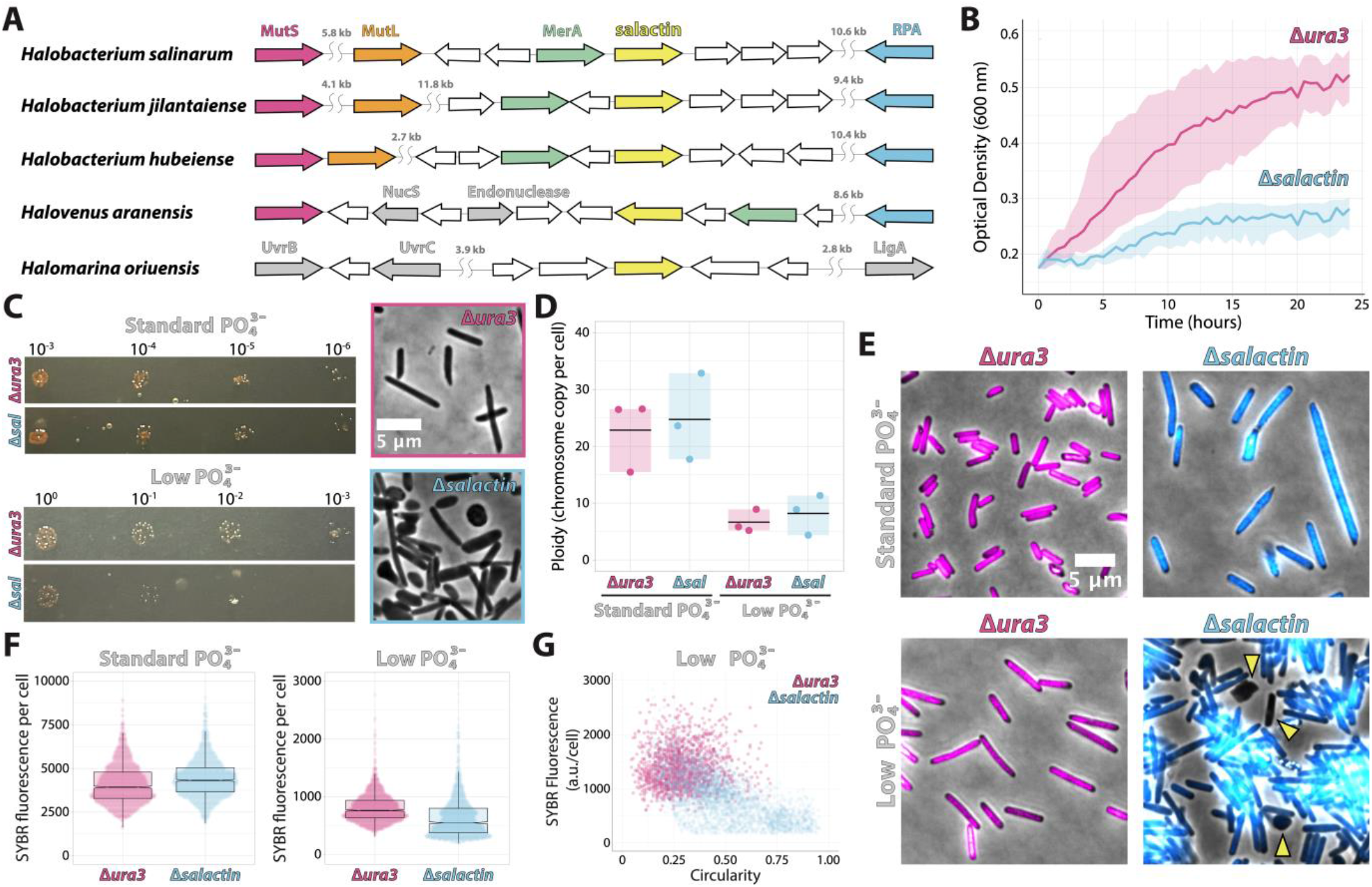
Cells lacking Δsalactin show defects in chromosomal partitioning and cell shape in low phosphate media. Throughout, pink is Δura3, and blue is Δsalactin. (A) Synteny analysis of salactin (VNG_RS00630; VNG0153C) gene in closely related organisms to H. salinarum. (B) Growth curve showing the difference in the growth of Δsalactin cells in low phosphate media. (C) Spot dilutions of Δura3 and Δsalactin cells grown on standard phosphate media and low phosphate media indicating Δsalactin cells have a reduced viability in low phosphate media relative to Δura3 cells (left). Representative phase images of Δura3 and Δsalactin cells from the spot dilution assay in low phosphate media (right). (D) The bulk chromosomal number per cell by qPCR in standard phosphate and low phosphate reveals that there is no statistical difference (p > 0.05) in the average chromosomal number per cell regardless of media condition. (E) Representative fluorescent images overlayed on the phase of Δura3 and Δsalactin cells stained with SYBR safe DNA stain in standard (left) and low phosphate media (right). Yellow arrowheads indicate cells lacking chromosomal material. (F) Quantification of the fluorescent intensity of SYBR safe stained Δura3 and Δsalactin cells in standard and low phosphate media. (p < 0.0001 for both). (G) SYBR fluorescence versus circularity of ura3 and Δsalactin cells in low phosphate media at stationary phase.

To test if Δ*salactin* cells showed a phenotype under decreased ploidy, we grew cells in a low phosphate media. Because Δ*ura3* cells failed to grow in a medium lacking any phosphate, we grew our cells in a media that combined a defined media lacking phosphate (Zerulla et al., 2014) with 1% of rich standard phosphate (CM) media to allow growth. In this growth condition, Δ*salactin* cells had a greatly decreased growth rate relative to Δ*ura3* cells, with Δ*salactin* cells considerably slowing down before one doubling (Fig 4B). Spot dilution assays indicated Δ*salactin* cells decreased growth rate in phosphate-limited media is likely due to a defect in viability: Δ*salactin* cells were indistinguishable from Δ*ura3* cells spotted on standard phosphate media but had at least two orders of reduced viability on low phosphate plates. In addition, when Δ*salactin* and Δ*ura3* cells were imaged directly from colonies on low phosphate spot dilution plates, there was a clear phenotypic difference in the shape of cells. Δ*ura3* cells were still consistently rod-shaped while Δ*salactin* cells had cells that became rounder and amorphous (Fig. 4C).

To understand if the drop in viability in Δ*salactin* cells was caused by a difference in ploidy compared to the Δ*ura3* parental strain, we performed qPCR on cultures of each strain. In standard phosphate media, Δ*ura3* and Δ*salactin* populations showed ploidy levels indistinguishable from each other (23±6.5 and 24±7.4 chromosomal copies, respectively), consistent to what was previously described (Breuert et al., 2006). In low phosphate media, we observed the chromosomal copy-number decrease approximately 3-fold for Δ*ura3* and Δ*salactin* cells (6.6±2.0 and 8.2±3.4 chromosomal copies, respectively) (Fig. 4D). Similar to the standard phosphate conditions, qPCR of Δ*ura3* and Δ*salactin* strains grown in low phosphate media also did not show any statistically significant difference in ploidy, revealing that, at the population level, the number of chromosomes per OD remains constant in both growth conditions in the presence or absence of Salactin (Fig. 4D). This indicates that Salactin is likely not involved in DNA replication or the degradation of DNA that occurs in response to phosphate depletion.

Next, we examined how the absence of Salactin affected the distribution of DNA within single cells, as well as their overall cell shape. We labeled the cellular DNA with SYBR safe DNA stain, and imaged cells with both phase contrast and widefield fluorescent microscopy. We performed these assays when cells were close to stationary phase, when cells have reduced ploidy (Breuert et al., 2006; Zerulla et al., 2014) as this might reveal greater phenotypic effects.

In standard phosphate media, while we could not observe a significant difference between the shape of Δ*ura3* and Δ*salactin* cells in exponential phase (Fig. 1C), Δ*salactin* cells were significantly larger than Δ*ura3* cells in stationary phase (Fig. 4E, S10A). Also, Δ*salactin* cells accumulated slightly more (7.7%) SYBR fluorescence relative to Δ*ura3* cells (p < 0.0001) (Fig. 4F, left panel). In contrast, comparing the SYBR intensities between cells in standard phosphate and low phosphate media revealed a clear downward shift for both Δ*ura3* and Δ*salactin* cells in low phosphate (Fig. 4F, right panel), in line with the reduced ploidy in our qPCR data when cells are shifted from standard to low phosphate media (Fig. 4D).

In low phosphate media, the amount of SYBR fluorescence in *Δsalactin* cells was far more heterogeneous relative to Δ*ura3* cells: some cells appeared to have little to no chromosomal material, while others appeared to have greatly increased DNA staining (Fig. 4E, S10B). Quantifying the amount of SYBR fluorescence per cell revealed a significant difference (p < 0.0001) between Δ*salactin* and Δ*ura3* cells, where the Δ*salactin* strain displayed a more bimodal distribution, with a new second peak appearing at the low end of the distribution (Fig. 4F, right panel). Δ*salactin* cells also showed shape defects in low phosphate media, with a large fraction of misshapen and amorphous cells, in contrast to the consistently rod-shaped Δ*ura3* cells (Fig. 4E). Interestingly, misshaped Δ*salactin* cells (higher circularity values) often correlated with reduced DNA staining (Fig. 4G). Given the average number of chromosomal copies per cell is the same between *ura3* and Δ*salactin* strains when assayed in bulk, these single cell results suggest that Salactin is required for correctly partitioning DNA between cells when ploidy is reduced, causing cells lacking Salactin to have less (or no) DNA, thereby making them unable to maintain their shape and overall viability.

## Discussion

Our studies revealed that, despite its homology to MreB, Salactin does not act as an MreB in *H. salinarum* cells. Rather, Salactin is a dynamically unstable polymer that grows and shrinks from the poles and appears to be involved in partitioning DNA between daughter cells when chromosomes become limiting. This represents the third actin homolog shown to exhibit dynamic instability (Garner et al., 2004; Petek et al., 2017), and the first characterization of the *in vivo* dynamics of any archaeal actin homolog.

Similar to other dynamically unstable polymers (like microtubules, ParM, Alp7A, and PhuZ) Salactin filaments grow from one end while the incorporated monomers remain immobile within the growing filaments (Desai & Mitchison, 1997; Garner et al., 2004; Sammak & Borisy, 1988). Diffraction limited Salactin filaments also appear to be composed of multiple filaments, similar to what was observed for ParM filaments in *Escherichia coli* (Campbell & Mullins, 2007; Salje & Löwe, 2008). Likewise, as required for other dynamically unstable actins, Salactin’s *in vitro* critical concentration in the non-hydrolyzed ATP state (∼0.3 µM) is lower than the ADP form (>10 µM), providing the energetic differential required for dynamic instability (Garner et al., 2004; Petek et al., 2017). The critical concentration that arises in hydrolyzable ATP (∼0.9 µM) likely reflects the relative proportions of growing and shrinking filaments in solution (Garner et al., 2004; Petek et al., 2017). Given that different salt concentrations can substantially alter the critical concentration of eukaryotic actin (Nakamura et al., 2011), our *in vitro* measurements of Salactin’s critical concentration conducted at much lower salt concentrations are likely not the same as those inside the cell and may explain why we were unable to recapitulate dynamic instability *in vitro*. However, even at our reduced *in vitro* salt concentrations, these measures suggest there exists a large difference in the ADP and non-hydrolyzable ATP critical concentrations as required for dynamic instability in the cell.

Cells lacking Salactin displayed substantial phenotypes only when cells were grown in low-phosphate media. *H. volcanii* has been shown to reduce its number of chromosomes from 30 down to as low as 2 in low phosphate media (Zerulla et al., 2014), and similarly, we also observe a reduction in *H. salinarum’s* ploidy and amount of DNA from 23 to 25 chromosomes down to 7 to 8 chromosomes in low phosphate media. Given Δ*salactin* cells show a DNA partitioning defect in low phosphate, it is likely that Salactin is part of a DNA segregation system that, similar to low copy plasmids, is required when the DNA copy number is not sufficiently high to be passed to both daughter cells by random chance. An inability to partition limiting chromosomes could create the apparent anucleate Δ*salactin* cells, which then would explain their defects in bulk growth, viability, and cell shape. Alternatively, given Salactin’s chromosomal proximity to multiple DNA repair enzymes in *H. salinarum* and its closest relatives (Fig. 4A), Salactin could also be involved in DNA repair and the SOS response.

Given all other known dynamically unstable filaments (microtubules (Kline-Smith & Walczak, 2004), ParM (Garner et al., 2004), Alp7A (Derman et al., 2009), and PhuZ (Erb et al., 2014) are involved in DNA partitioning, our results suggest that Salactin filaments could partition archaeal DNA by their growth or catastrophe, pulling from some “kinetochore-like” region on the archaeal chromosome. Because we currently lack the tools to label discrete loci on chromosomes, we were unable to verify our model, but future studies that simultaneously image Salactin filaments and discrete chromosomal loci could test this model.

## Materials and Methods

### Strains, plasmids, and primers

Halobacterium salinarum NRC-1 (ATCC 700922) was the wild-type strain used in this study. Table S4, S5, S6 lists strains, plasmids, and primers used in this study. For more detailed plasmid construction, see the supplementary methods. The pRrpa plasmids were created from a modified version of the pMTFChis (Darnell et al., 2020), where the original promoter, P_fdx_, is replaced with another promoter, P_rpa_. Plasmid constructs for the transformation of *H. salinarum* were generated by isothermal Gibson assembly (Gibson, 2011) of: 1) the PCR fragments (amplified by KAPA Biosystems DNA polymerase [VWR] and gel extracted); and 2) the linear plasmid. Plasmids were propagated in *Escherichia coli* DH5. Proteins were tagged with fluorescent proteins as C-terminal fusions using a 15 amino acid linker (LEGSGQGPGSGQGSG). Fluorescent protein sequences (msfGFP and HaloTag) were obtained from Dion and colleagues (Dion et al., 2019). Plasmids were verified by Sanger sequencing of the locus in the plasmid. Plasmids were transformed into *H. salinarum* using a polyethylene glycol 600 spheroplast protocol (Cline et al., 2011; Dyall-Smith, 2009) and selected using mevinolin. The protocol used for protein overexpression and purification using the his6SUMO (Malakhov et al., 2004) construct (pSUMO) is described by Stoddard and colleagues (Stoddard et al., 2020).

### Media and growth conditions

*H. salinarum* strains were routinely grown, unless otherwise specified, using a nutrient-rich medium, CM (Complete Media) medium (250 g/liter NaCl [Fisher Scientific]; 20 g/liter MgSO_4_·7H_2_O [Fisher Scientific]; 3 g/liter trisodium citrate [Fisher Scientific]; 2 g/liter KCl [Fisher Scientific]; 10 g/liter bacteriological peptone [Oxoid]; pH 6.8). Media were supplemented with 50 g/ml uracil (Sigma) to complement the uracil auxotrophy of the *Δura3* background. All growth was performed at 42°C in a roller drum. Self-replicating *H. salinarum* plasmids were maintained using 1 g/ml mevinolin in liquid culture. Cells were grown at 37°C during live-cell microscopy. *E. coli* was grown in an LB medium with carbenicillin (50 g/ml; Sigma) to maintain plasmids. For *H. salinarum* growth in low phosphate, a phosphate-free media, which was adapted from Zerulla and colleagues was prepared (Zerulla et al., 2014) (10 mM NH_4_Cl; 0.5% Glucose; 25 mM CaCl_2_; 10 mM Trisodium Citrate·2H_2_O; 25 mM KCl; 100 mM MgSO_4_·7H_2_O; 4.2M NaCl; Trace Metal Solution; Vitamin Solution; pH 6.8). Trace metal and vitamin solutions were prepared as described by de Silva and colleagues (de Silva et al., 2021). The low phosphate condition consisted of the phosphate-free medium mixed with rich CM 99:1 prior to *H. salinarum* culturing.

### Data Visualization

Data was plotted with GraphPad Prism (Prism 9 for macOS version 9.3.1 (www.graphpad.com), PlotsOfData https://huygens.science.uva.nl/PlotsOfData/ (Postma & Goedhart, 2019), and PlotTwist https://huygens.science.uva.nl/PlotTwist (Goedhart, 2020).

### Genomic Neighborhood Analysis

Synteny of *salactin* across different haloarchaeal genomes was performed using Syntax (https://archaea.i2bc.paris-saclay.fr/SyntTax/) (Oberto, 2013). Salactin protein sequence from *H. salinarum* was used as bait in search against deposited haloarchaeal genomes with a 20% minimal normalized genomic BLAST score.

### Whole genome sequencing

*H. salinarum* were grown to mid/late-logarithmic phase (optical density at 600nm (OD_600_) ∼0.7) and 1.5mL was pelleted by centrifugation and stored at -20°C until processed. Cells were lysed in ddH_2_O and DNA was extracted using a phenol-chloroform method in phase lock gel tubes and ethanol precipitated and washed. DNA pellet was resuspended in TE buffer and quantified using a NanoDrop. Samples were submitted to the Bauer Core Facility in the FAS Division of Science at Harvard University for Nextera XT ¼ volume queued library prep and an Illumina MiSeq v2 run for whole genome sequencing. Sequences were trimmed by BBDuk trimmer prior to whole genome assembly using Geneious de novo assembly of paired-end plugin. The results of this analysis can be found in Table S1.

### Phylogenetic analysis

We performed iterative HMM searching to retrieve Salactin homologs from a database of 700 representatively sampled archaeal and bacterial proteomes (Moody et al., 2022). Beginning with sequence record WP_010902067.1, we retrieved all phmmer hits with E < 1e-7. These sequences were aligned, a new HMM was built, and additional homologs were retrieved from the database. This search returned an expanded set of hits, including other sequences in Methanotecta, Euryarchaeota more broadly, and more distant members of the actin fold family (included in the expanded tree in Fig. S1). The focal phylogeny (Fig. 1A) includes sequences from Haloarchaea and other Methanotecta, which branched closest to the initially identified Salactin sequences in the expanded phylogeny (Fig. S1). To place Salactin and its orthologs in Methanotecta in the broader context of the actin superfamily phylogeny, we also inferred a phylogeny including homologs of MreB (PF06723) more broadly. All sequences were aligned with mafft (l-ins-I mode) (Katoh & Standley, 2013) and trimmed using BMGE with a blosum30 matrix (Criscuolo & Gribaldo, 2010) prior to tree inference. Phylogenies were inferred under the LG+C20+F+G model in IQ-TREE 2 (Minh et al., 2020) with 10,000 ultrafast bootstraps (Hoang et al., 2018). This substitution model was the best fitting model for the focal analysis according to the BIC criterion.

### Growth curves

i. Rich media: Liquid cultures were grown in CM (3mL) from single colonies until saturation. Cells were then diluted to an OD_600_ of 0.025, representing time 0 for all growth curves. OD measurements were taken by hand every 6-12 hours throughout the growth curve.
ii. Low phosphate media: Cultures were grown in rich media (CM) as described above, then centrifuged at 4,000g for 5 minutes and washed three times with phosphate-free media. After washing, cells were resuspended to an initial OD_600nm_ of 0.2 in low phosphate media. 200 µl of each sample (and fresh media as a blank) were then transferred to a 96-well plate to be read in a BioTek EPOCH2 plate reader. Growth curves were obtained by taking OD_600_ readings every 30 minutes for 24 hours, under continuous orbital shaking (425 cpm) at 42°C. Data was plotted using PlotTwist (https://huygens.science.uva.nl/PlotTwist) (Goedhart, 2020).

### Motility assay and diameter measurements

Liquid cultures were grown in CM from single colonies until saturation. Cells were then diluted to an OD_600_ of 0.025 in CM and grown to early log phase (OD_600_ 0.1-0.3). The OD_600_ of these cells was normalized, and then the cells were stabbed onto a 0.3% 1:10 CM [same composition as CM but 1/10 Oxoid peptone] agar plate for the motility assay. The diameter was measured by hand on day 6 of incubation at 42°C in a thick plastic bag to control the loss of humidity.

### Spot dilution viability assay

Δ*ura3* and Δ*salactin* cells were grown in rich media (CM) until exponential phase (OD_600nm_ = 0.5) and then serially diluted in fresh rich media. 2 µl of each diluted cultures were spotted onto rich CM and low-phosphate agar plates and incubated at 42°C.

### Ploidy determination by qPCR

Standard curves were generated with isolated gDNA from cultures grown in the spot dilution assay described above. Genomic DNA was isolated using the Quick-DNA Fungal/Bacterial Miniprep Kit (Zymo Research). Concentrations were determined by NanoDrop (Thermo Scientific 2000) and each sample was diluted to 0.4 ng/µl. Samples were serially diluted so their final working concentrations fell within the range of the generated standard curve. The standard curve was created using a 270bp PCR product (Breuert et al., 2006). oJM220 and oJM221 primers were designed using NCBI’s PCR primer design tool (https://www.ncbi.nlm.nih.gov/tools/primer-blast/). A standard PCR reaction was performed at 95°C for 2 minutes (denature), followed by 35 cycles of 15 seconds at 95°C (denature), 15 seconds at 55°C (annealing), and 20 seconds at 68°C (extension). The 270bp amplicon was purified from an agarose gel using the Zymoclean Gel DNA Recovery Kit (Zymo Research). qPCR was performed on a 10-fold dilution series of the purified product from 0.1ng to 1×10^−6^ng of template. Template concentration versus Cq values was plotted and a linear line was fit to the data. The equation of this line was used to determine DNA amounts in the ploidy experiments. The qPCR reactions were performed using a clear, 96-well PCR plate (Olympus Plastics). To each well 7.5ul of a master mix was added, consisting of SYBR Green/dNTPs/Taq polymerase/and primers at 0.25 µM. A melt-curve was then determined by heating the samples from 65°C to 95°C in 0.5°C steps. The qPCR was performed in the CFX96 Real-Time System (Bio-Rad). Data analysis was performed and Cq values were determined using the CFX Maestro software (Bio-Rad). DNA was extracted from a total of three biological replicates and all samples were run in triplicate with water acting as a template control. Calculated Cq values were compared to a standard curve and divided by the colony formation unit (cfu) from initial cultures to determine copy-number of the main chromosome per cell in each sample.

### Imaging

Unless otherwise noted, strains were grown in liquid culture (3mL CM) from single colonies until saturation. Cells were then diluted to OD_600_ 0.1 and grown to exponential phase (OD_600_ 0.5-0.8) before the start of all imaging experiments,

i. Cell size and shape measurements. Strains: *Δura3, Δsalactin*. For imaging, 5uL culture aliquots were immobilized on No. 1.5 cover glass under a CM agar pad (0.3% w/v). Phase-contrast images (100ms exposure) were collected on a Nikon TI microscope equipped with a 6.45-µm-pixel Andor Clara camera and a Nikon x 100 numerical aperture (NA) 1.4 objective. The phase-contrast images were segmented manually, and size and shape measurements were obtained using Fiji’s measure function. The difference between the two groups was analyzed using the Mann-Whitney test found in the t tests (and nonparametric tests) analysis on Prism 9 for macOS version 9.3.1 (www.graphpad.com) for all statistical analyses unless otherwise stated.
ii. Imaging of dynamics. Strains: WT + *prpa-salactin-msfGFP* and *prpa-salactin-halotag, Δsalactin* + *prpa-salactin-msfGFP* and *prpa-salactin-halotag*. For imaging, 5uL culture aliquots were immobilized on No. 1.5 cover glass under a CM agar pad (0.3% w/v). Cells were imaged in a Nikon Eclipse Ti microscope with a 6.5-m pixel ORCA-Flash4.0 V2 sCMOS Hamamatsu camera and a Nikon 60 NA 1.4 phase-contrast objective for comparing dynamics between conditions and a 100 NA 1.45 phase-contrast objective for dynamic phenotypes. Fluorescence excitation was achieved using a MLC4008 laser launch (Agilent), with a 488nm laser used for msfGFP imaging and a 561nm laser used for imaging of JF549 conjugated to the HaloTag. Using fluorescence HiLo microscopy, images were captured every 30 seconds for 30 minutes to 1 hour at 40% laser power. Exposure times for fluorescence were 100ms and 200ms for the 488nm and 561nm laser respectively. The polymerization rates, length, and time for catastrophe were measured by kymograph analysis. Kymographs were created from time lapses of fluorescently labeled Salactin filaments by manually drawing ROIs along the long axis of the cells in Fiji or along the filament (whichever created clearer kymographs). Regions of these kymographs containing right triangles represented dynamic instability and were measured manually in Fiji for the dynamic measurements. For the first exponential distribution analysis, time for catastrophe is plotted as a relative frequency histogram with a bin size of 150 and fitted to a one-phase decay exponential in the “non-linear regression (curve fit) analysis” in Prism 9 (www.graphpad.com). For the second fitting method, the logarithm of frequency is used to create a linear fit using the “simple linear regression analysis” in Prism 9 (www.graphpad.com).
iii. Single-molecule. Strains: WT + *prpa-salactin-halotag*. To decrease autofluorescence, liquid cultures were grown in HS-Ca media, which was modified from the Hv-Ca media (Allers et al., 2004)(Allers et al., 2004)(Allers et al., 2004)(Allers et al., 2004)(Allers et al., 2004)(Allers et al., 2004)(Allers et al., 2004) (3mL) (Ingredients: 25% BSW (240g/L NaCl [ChemSupply: SA046], 30g/L MgCl_2_·6H_2_O [Sigma: M2393], 35gL MgSO_4_·7H_2_O [Sigma: V800245], 7g/L KCl [Sigma: V800245], 20mL 1M Tris-HCl pH7.4 [ChemSupply: TA034]), 0.5% w/v Casamino acids (5g/L [Oxoid:LP0041]), from single colonies until saturation. Cells were then diluted to OD_600_ 0.1 to grow to late exponential (OD_600_ 0.75-0.85). Cells were conjugated with a mixture of JF dyes (Grimm et al., 2015) to perform single-molecule and whole-cell labeling to verify the localization of the single molecules. JF dyes were added to the growth media 15 minutes before imaging. The ratio of the dyes was 1:20 JF549:JF505. 1.25nM JF549 was used and 25nM JF505 was used. For imaging, 5uL culture aliquots were immobilized in Matek dishes under an HS-Ca agar pad (0.3% w/v) to optimize signal-to-noise. Cells were imaged in a Nikon Eclipse Ti microscope with a 6.5-m pixel ORCA-Flash4.0 V2 sCMOS Hamamatsu camera and a Nikon 100 NA 1.45 phase-contrast objective. Fluorescent images were obtained using a MLC4008 laser launch (Agilent), with a 488nm laser for JF505 imaging and a 561nm laser for the JF549 imaging. Using HILO microscopy, fluorescent images were captured every 5-10 seconds for 30 minutes to 1 hour. Exposure times for fluorescence were 1s (561nm laser) and 250ms (488nm laser).
iv. DNA labeling in live cells. Cells were grown in rich media until exponential phase (OD_600_ ∼0.5), then centrifuged at 4,000g for 5 minutes and washed three times with phosphate-free media. After washing, cells were resuspended to an initial OD_600_ of 0.2 in rich or low-phosphate media and grown to saturation. Live-cell DNA labeling was done by adding SYBR Safe DNA Gel Stain (Invitrogen) to a final 10^6^-fold dilution. Cells were then incubated for 5 minutes at room temperature and promptly imaged. The difference between the DNA labeling of the two groups was analyzed using the Mann-Whitney test found in the t tests (and nonparametric tests) analysis on Prism 9 for macOS version 9.3.1 (www.graphpad.com)
v. Visualization of filaments in TIRF. 7µM of Salactin was mixed with 1/20^th^ of stained Salactin-GSKCK. To visualize Salactin fluorescently, Salactin-GSKCK was conjugated with cy3B mono maleimide. To conjugate the protein to the dye, the protein was pre-reduced with 5mM TCEP for 30min. TCEP was removed with a desalting column (NAP-5), and 5x excess dye was added and incubated for 10min on ice. The reaction was quenched by adding 10mM DTT. The protein was hard spun on a TLA100 rotor at 90K rpm for 20min. The supernatant was loaded on a NAP-5 desalting column, and concentration was measured based on the A_280_. Polymerization reactions were initiated upon adding 3mM MgATP, or MgAMPPNP, with 17.6% PEG8000 to protein and incubated at 37C for 3hr. Without any PEG, filaments could not be visualized under TIRF. For imaging, 5uL of protein solution was immobilized between two No. 1.5 cover glasses (22×60mm base and 18×18mm cover) and visualized using TIRF microscopy on a Nikon Eclipse Ti microscope with a 6.5-m pixel ORCA-Flash4.0 V2 sCMOS Hamamatsu camera and a Nikon 100 NA 1.45 phase-contrast objective. Fluorescent images were obtained using a MLC4008 laser launch (Agilent) with a 561nm laser to image the cy3B dye. Images were captured with a 200ms exposure time at 30% laser power.

### Protein Purification

BL21 (DE3) Rosetta containing the his6SUMO fusion plasmid was grown to OD_600_ ∼0.6 and induced for four hours with 0.4mM IPTG at 37°C. Cell pellets were resuspended in I0 buffer (Ingredients: 50mM Tris, 300mM KCl, 1mM MgCl_2_, 10% glycerol. Added before use 0.5mM TCEP and 0.2mM ATP) and stored at -80°C until use. Cells were lysed using a Misonix Sonicator. His6SUMO fusion products were then purified using a 5mL HisTrap HP (GE Healthcare) on an AKTA pure with stepwise imidazole·HCl increases from 15mM, 35mM, 50mM, 65mM, to 80mM and a final gradient to 500mM imidazole·HCl (I500). The His6SUMO tag was cleaved off using the Ulp1 protease (Malakhov et al., 2004) during dialysis of the protein back into I0. Using gravity, Proteins were then run through a 4mL bed of HisPur Ni-NTA resin (Thermo Scientific) with a wash step of I0 buffer (where the desired cleaved protein will come out) and a I500 buffer elution step. Proteins were further purified using a 5mL HiTrap Q FF (GE Healthcare) on an AKTA pure. Protein was dialyzed into starting buffer for anion exchange (Ingredients: 20mM Tris pH 8, 30mM KCl, 1mM MgCl_2_. Added before use 0.5mM TCEP and 0.2mM ATP) and bound to the HiTrap Q. The column was washed with starting buffer (5CV), 20% of elution buffer (5CV) (Ingredients: 20mM Tris pH8, 1M KCl, and 1mM MgCl2. Added before use 0.5mM TCEP and 0.2mM ATP), eluted using a gradient from 20% elution buffer to 65% elution buffer, and washed with 100% elution buffer. Fractions with protein were pooled and concentrated using a 10k MWCO PES Pierce Protein Concentrator (Thermo Scientific). Proteins were buffer exchanged using a prepacked PD-10 desalting column into HP-buffer (2.9M KCl, 5mM MgCl2, 10mM HEPES (pH 7), and 0.2mM EGTA. Added before use: 0.2mM ATP and 0.5mM TCEP). The protein concentration was determined using a Pierce BCA Protein Assay kit [ThermoScientific] due to the low A_280_ signal from the absence of tryptophan in the protein. 25% glycerol was added to the protein, and aliquots were snap-frozen in liquid nitrogen and stored at -80°C until needed.

### Malachite Green Assay

Malachite Green Assays were performed using a malachite green phosphate assay kit [MAK307-1KT: Sigma-Aldrich]. Polymerization reactions were started by mixing at least 5.5µM protein (unless otherwise stated) with 0.2mM ATP and heated to 37C for varying times. The reaction was stopped by adding 25µM EDTA and 75µM sulfuric acid. The reaction was spun down for 5min at 16g, the supernatant was taken, and phosphate was measured using the malachite green assay kit.

### Pelleting Assay

Unless otherwise noted, polymerization reactions were done in HP buffer and contained 2.5 mM MgATP or MgAMPPNP. When testing other nucleotides, 2.5mM of the nucleotide is used. Protein was exchanged into HP buffer without ATP and initiated upon adding MgATP or MgAMPPNP. Polymerization reactions were run for 3hr at 37C. Polymerization reactions were spun in a TLA100 (Beckman) at 436k g for 30 minutes at 37°C. Supernatants were removed and added to an equal volume of 2xSDS-Buffer. Pellets were resuspended by heating at 65 °C in 2 volumes of 1xSDS-Buffer. Fractions were subjected to SDS-PAGE and stained with SYPRO Orange [ThermoFisher]. The gel was visualized on a c200 Azure gel imaging station, with the EPI Blue LED, with a 470nm wavelength, and band intensities were quantified in ImageJ (Schneider et al., 2012). Note that more qualitative data, like the pelleting comparison for different nucleotides, were taken on a blue box using an iPhone camera.

### Fluorimeter experiments

Protein was exchanged into HP buffer without ATP and initiated upon adding MgATP or MgAMPPNP. Light scattering polymerization reactions were initiated by mixing protein with 17.6% PEG8000 and 3mM MgATP or 2mM MgAMPPNP and incubated at 37C for 3hr. Without any PEG, there would be no signal of polymers forming after 3hr of incubation even though pelleting indicates polymerization occurs. All light scattering experiments were endpoint assays at 3hr. The 90-degree scattering of the solution at 315nm was measured using a Fluorolog-3 (Horiba).

## Supporting information

Excel file with WGS

Treefile #1

Treefile #2

## Acknowledgments

JZ was supported by a grant 203276/l/16/Z from the Wellcome trust, and support from the NSF-Simons Center for Mathematical and Statistical Analysis of Biology at Harvard (award #1764269). AKS acknowledges support through NSF-MCB CAREER award (1651117). T.A.W and E.R.R.M acknowledges support through the John Templeton Foundation (62220). The opinions expressed in this publication are those of the authors and do not necessarily reflect the views of the John Templeton Foundation. AB-F and JM are supported by the Moore-Simons Project on the Origin of the Eukaryotic Cell award (GR404060) and the NSF MRSEC Bioinspired Soft Materials Award (DMR-2011846). AB-F is a Pew Scholar in the Biomedical Sciences, supported by The Pew Charitable Trusts.

We acknowledge the support the Bauer Core Facility at Harvard University for their help with sequencing. We also thank the Baliga lab for providing the Δ*salactin* strain. We also thank Patrick Stoddard and Elizabeth May for all the support with the biochemical experiments as well as Paul Dieterle for the discussions on catastrophe models. We thank Simonetta Gribaldo and Nika Pende for insightful comments in our manuscript. We thank the MBL Woods Hole Physiology course for providing a space for breakthroughs and discoveries in this project.

