## Supplemental text 1 for "Salactin, a dynamically unstable actin homolog in Haloarchaea"

### SUPPLEMENTARY FIGURES

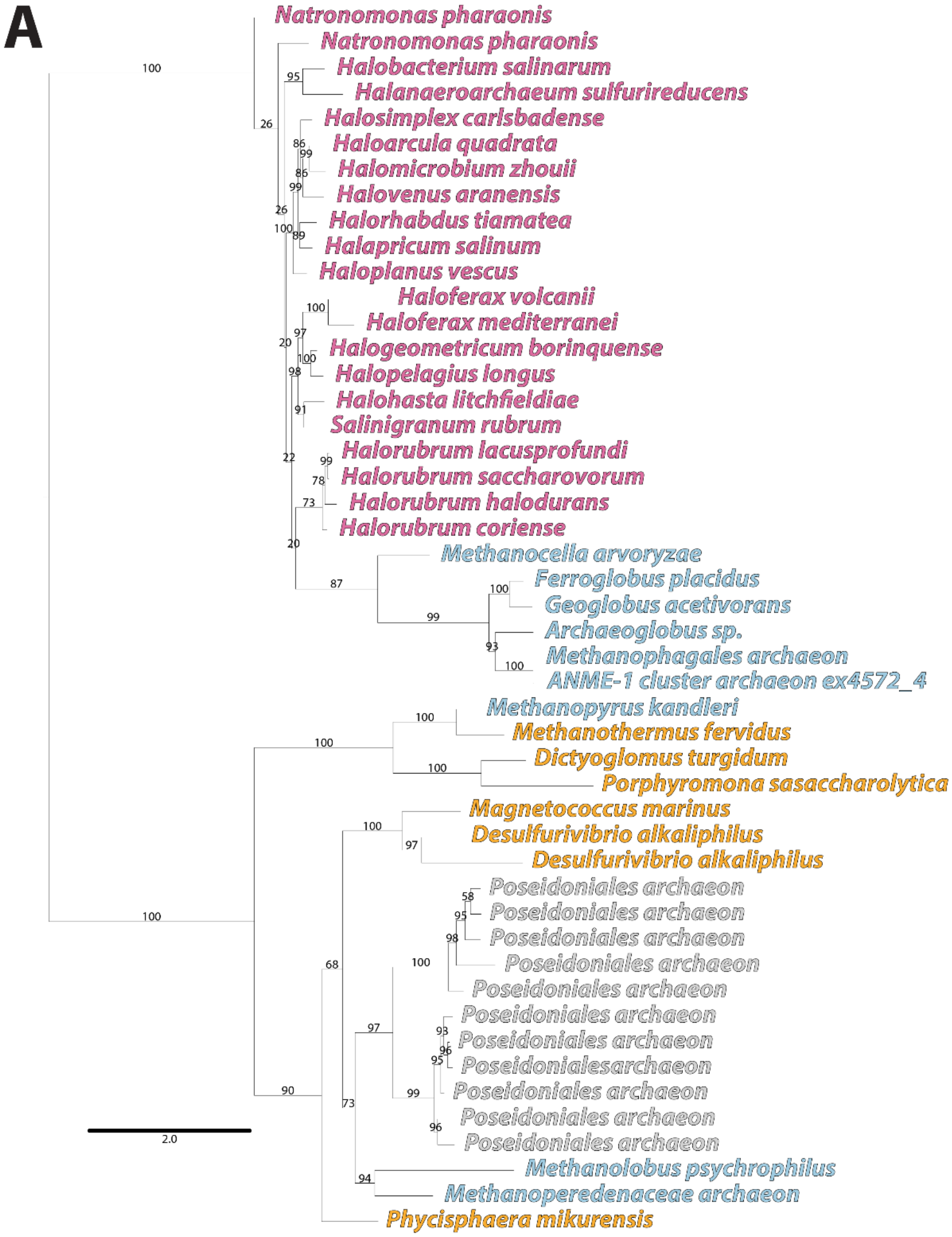


**Supplemental Figure S1.** Phylogenetic trees (A) Zoomed out tree of the representative species in the Euryarchaeota phylum with a closely related homolog of Salactin. The numbers on branches are the ultrafast bootstrap support and represent how reliable the branch is, where 100% is well supported, and greater than 95% is strongly supported. (B) Tree of Salactin in the context of other actin homologs (see separate pdf file). Salactin orthologs are pink, FtsAs are orange, MreBs are blue, Hsp70/DnaKs are green, and EutJs are grey. Note that this tree is drawn as rooted between Hsp70/DnaK and MreB, but this is for visualization purposes only as the deep relationships between Hsp70/DnaK, MreB and FtsA are not currently known with certainty (Stoddard et al., 2017).

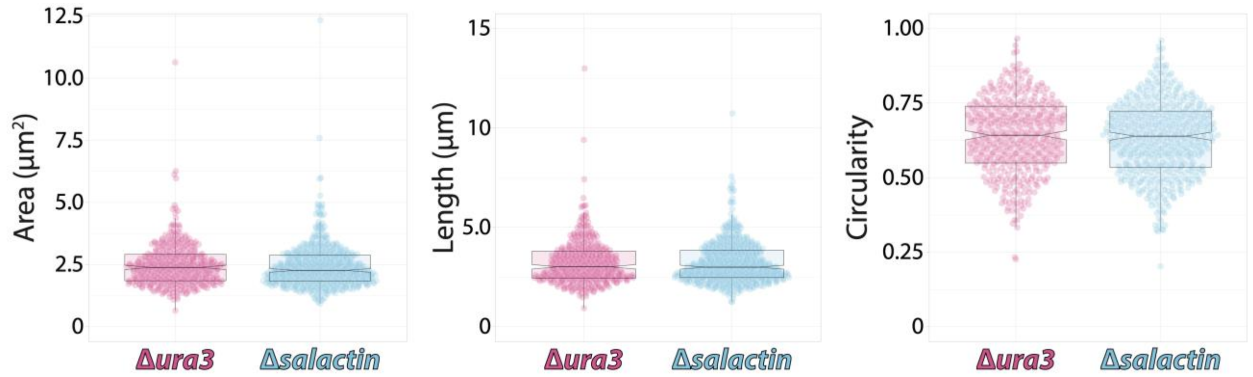


**Supplemental Figure S2.** Box plots showing no statistical differences between Δ*salactin* and Δ*ura3* cells in their (*left*) area (p-value = 0.21), (*mid*) length (p-value = 0.98), or (*right*) circularity (p-value = 0.34).

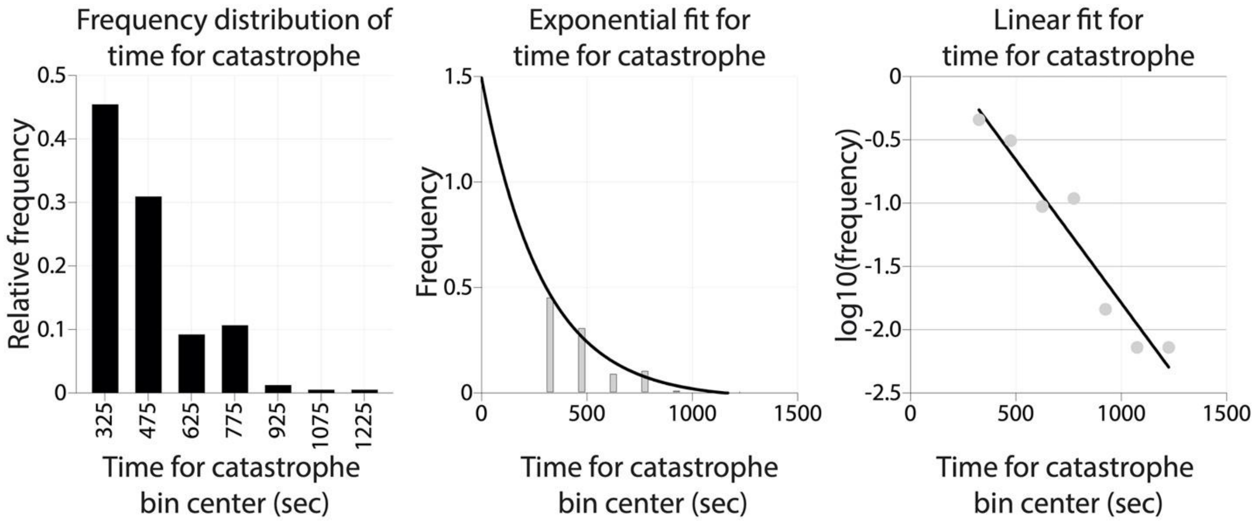


**Supplemental Figure S3.** Catastrophe time distribution analysis. Time for catastrophe plotted as a relative frequency bar graph with 150 bin sizes (*left*) and fitted to a one-phase decay exponential in the "non-linear regression (curve fit) analysis" routine in Prism 9 for macOS version 9.3.1 (www.graphpad.com) (*mid*). R-squared value = 0.9643. A second method of fitting done by calculating the logarithm of the frequency and creating a linear fit using the "simple linear regression analysis" routine in Prism 9 for macOS version 9.3.1 (www.graphpad.com) (*right*). R-squared value = 0.9335.

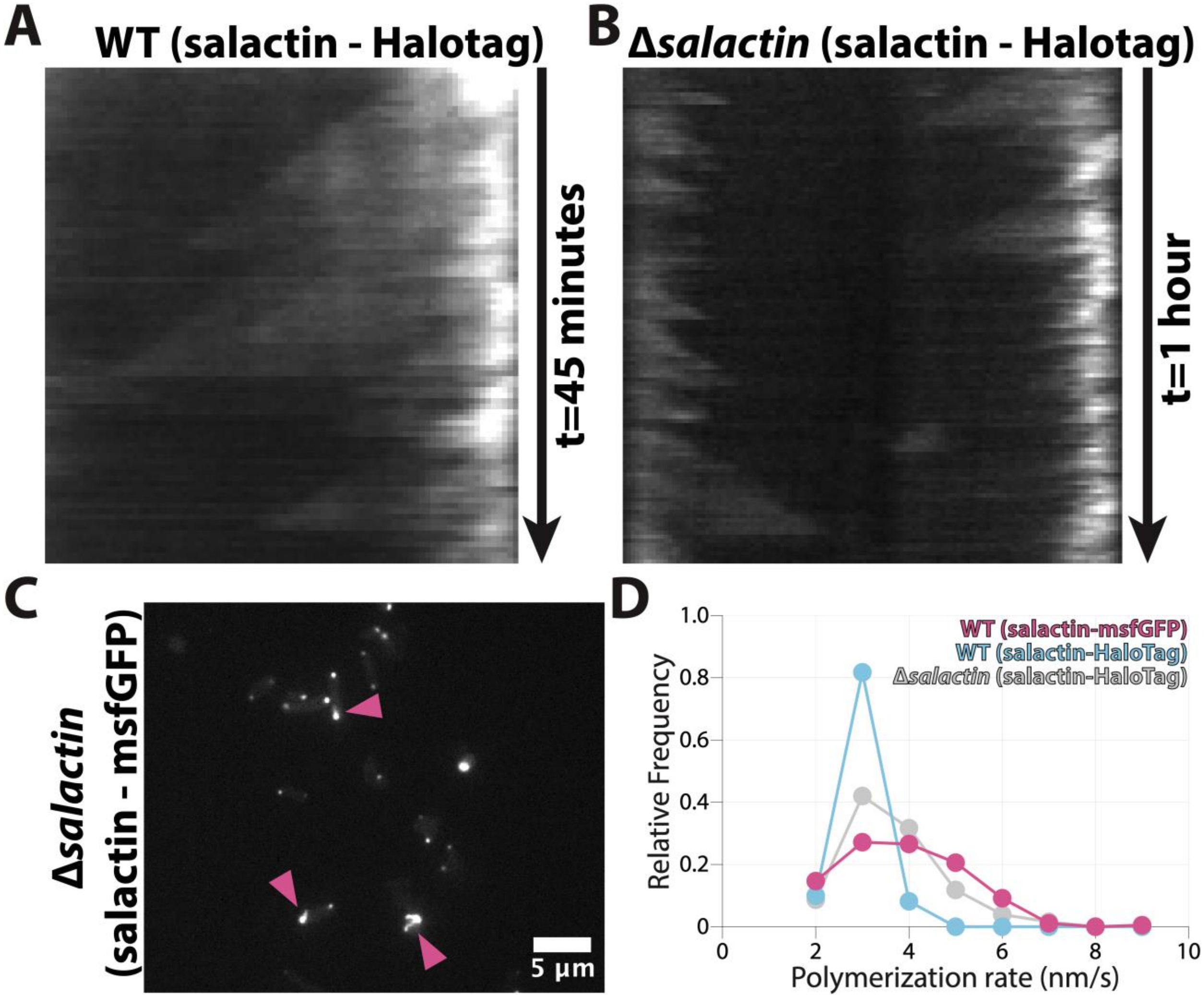


**Supplemental Figure S4.** Dynamics of Salactin fused to different tags. (A) Kymograph of Salactin-HaloTag (strain hsJZ86) also expressing the native *salactin* copy. (B) Kymograph of Salactin-HaloTag expressed in Δ*salactin* cells (strain hsJZ106). (C) Fluorescent images of Salactin-msfGFP expressed in Δ*salactin* cells (strain hsJZ95). The pink arrow points toward the appearance of filaments. (D) Histogram of the relative frequencies for polymerization rates comparing wild type + Salactin-msfGFP (strain hsJZ52), wild type + Salactin-Halotag (strain hsJZ86), Δ*salactin* + Salactin-HaloTag (strain hsJZ106). The wild type + Salactin-HaloTag has a significantly (p-value <0.0001) slower polymerization rate, while the other two have comparable polymerization rates.

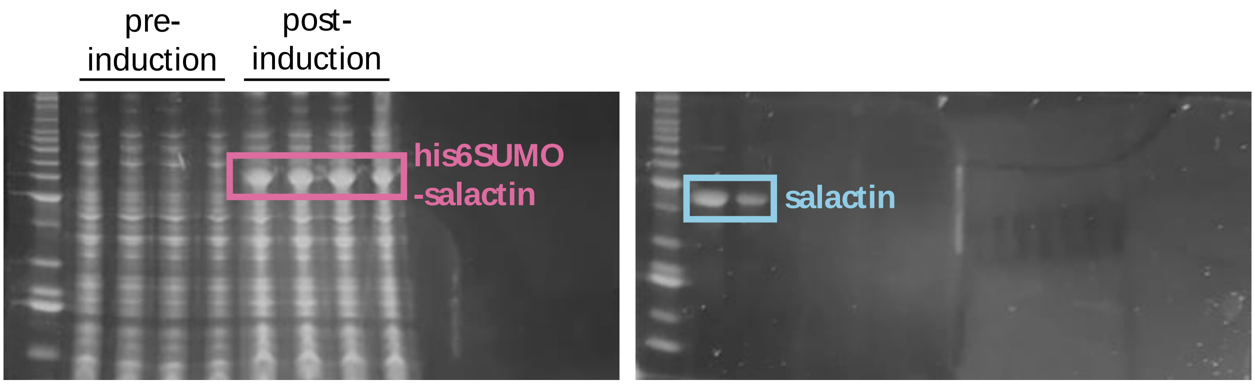


**Supplemental Figure S5.** SDS-PAGE gels of the purification protocol at the beginning (induction) and end (purified protein). Pre- and post-induction of Salactin expressed in *Escherichia coli* (*left*). Final purified protein used for *in vitro* assays (*right*). All gels are stained with SYPRO Orange.

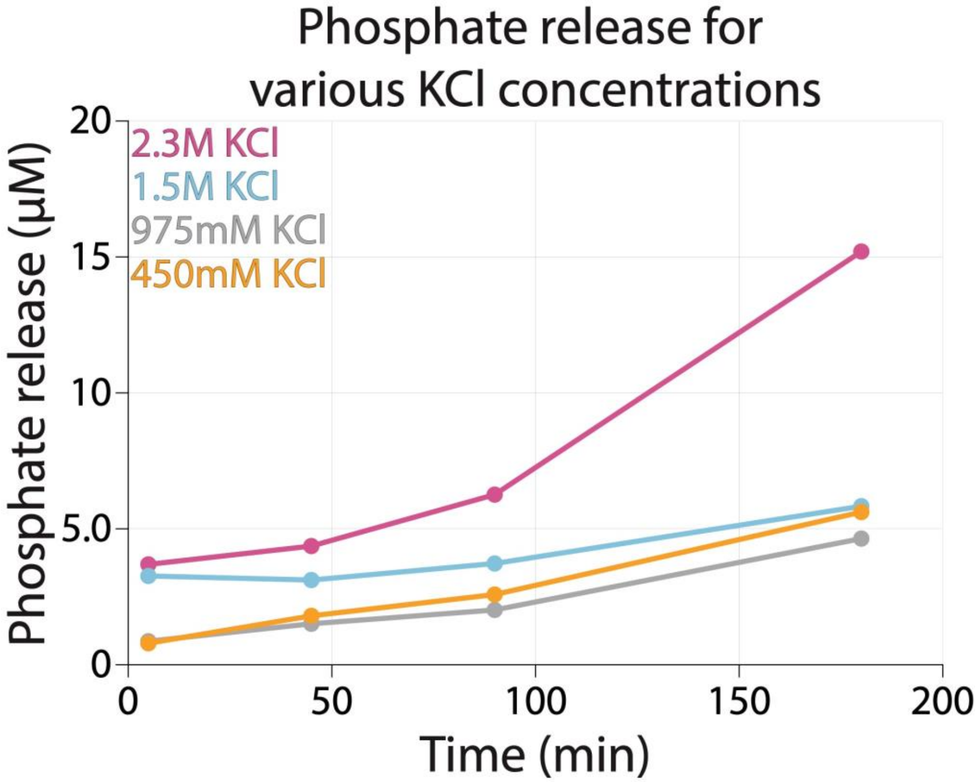


**Supplemental Figure S6.** Malachite Green Assay using 4 μM Salactin in different salt conditions (450 mM, 975 mM, 1.5 M, 2.29 M KCl). As noted in the main text, the higher ATPase activity suggests polymerization is favored at higher salt concentrations.

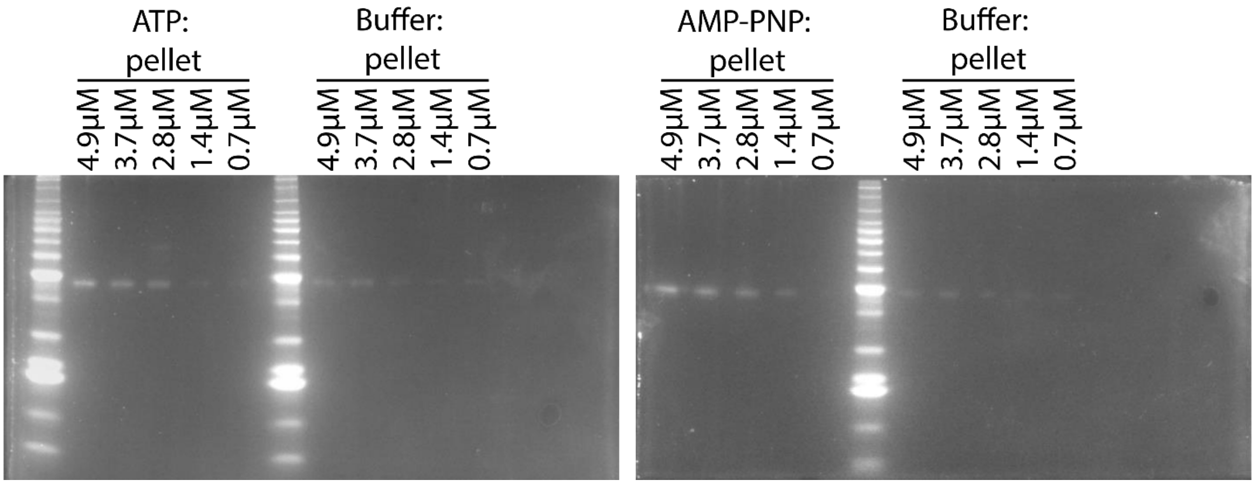


**Supplemental Figure S7.** Representative pelleting SDS-PAGE gels of Salactin in ATP (*left*) and AMPPNP (*right*) across different protein concentrations. All gels are stained with SYPRO Orange.

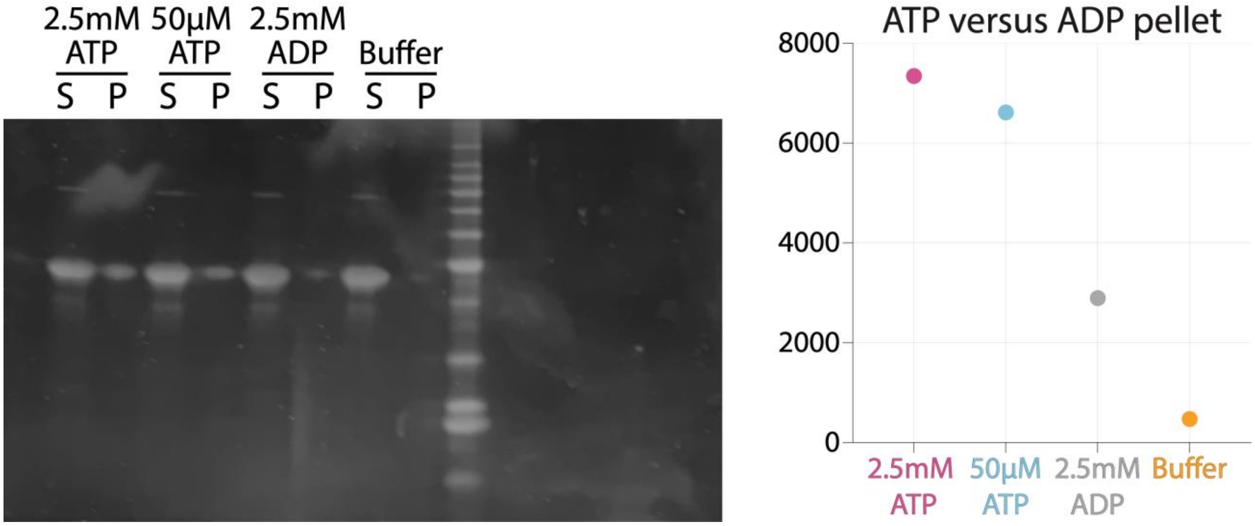


**Supplemental Figure S8.** Pelleting of Salactin in ATP compared to ADP. SDS-Page gel of 10 µM Salactin in 2.5 mM ATP, 50 µM ATP, 2.5 mM ADP, and HP-buffer (*left*). Measured SYPRO Orange intensity of the four different conditions (*right*). All gels are stained with SYPRO Orange. S = supernatant, P = pellet.

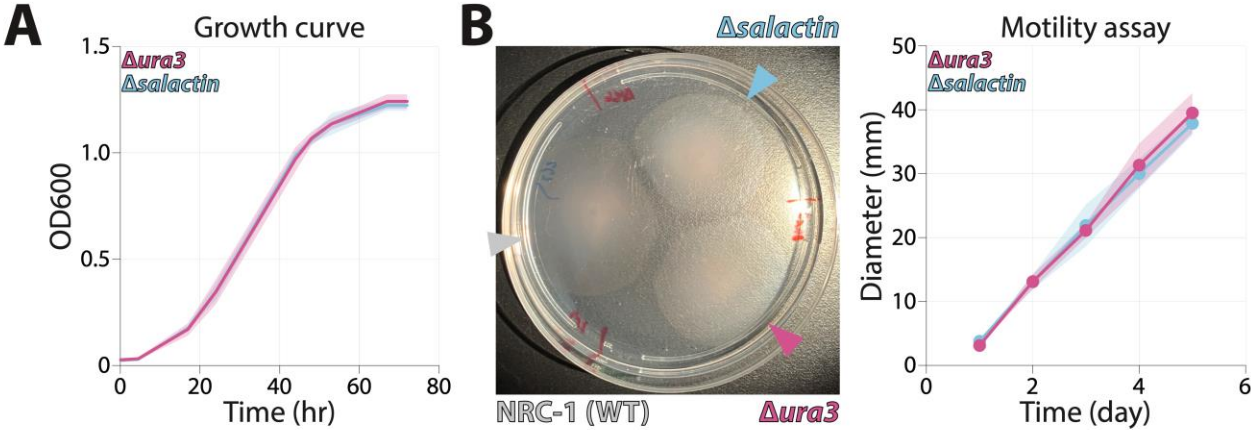


**Supplemental Figure S9**. (A) Growth curves of Δ*ura3* and Δ*salactin* cells in rich media. (B) Representative image of Δ*ura3* and Δ*salactin* on an agar motility plate (10% CM, 0.3% agar) 5 days after inoculation *(left*). Δ*ura3* is in the bottom right corner, Δ*salactin* is in the upper right corner of the plate, and NRC-1 (wild-type cells) are on the left. Quantitation of 9 plates across 5 days indicated no motility defect in the Δ*salactin* strain relative to the Δ*ura3* strain (p-value >0.05 for each of the five days) (*right*).

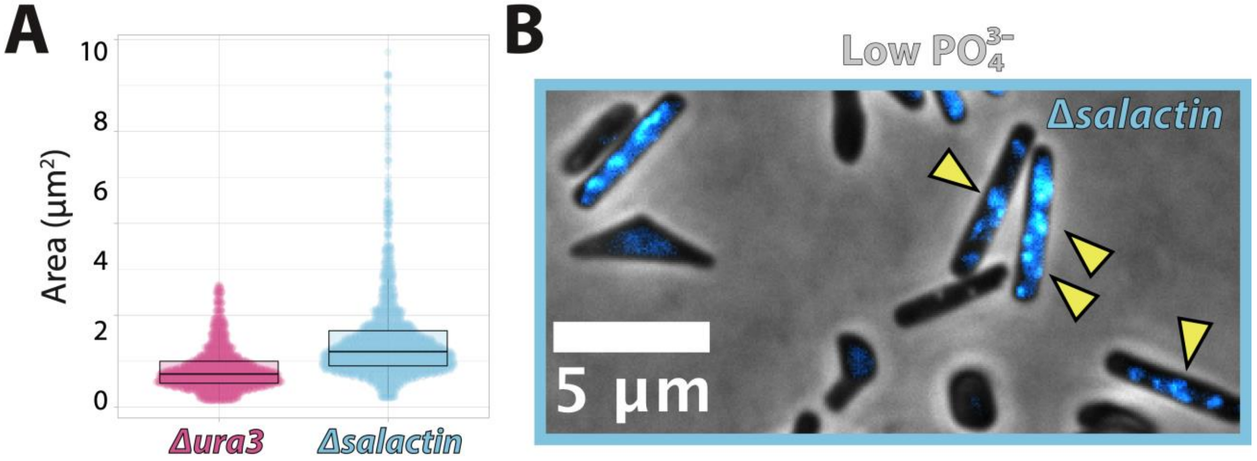


**Supplemental Figure S10**. (A) Area of Δ*ura3* and Δ*salactin* cells in standard phosphate media at stationary phase. (B) Zoomed in image of Δ*salactin* cells in low phosphate demonstrating that cells now show clear foci of DNA as indicated by the yellow arrowheads.

### SUPPLEMENTARY MOVIES

**SM1.** Video of Salactin-msfGFP expressed on top of the native copy, visualized by Near-TIRF fluorescent microscopy. Images were taken with a 488nm laser every 30 seconds for 1 hour. Video is 600x actual speed. Scale bar = 4 μm. Movie corresponds to the montage in Figure 2A.

**SM2.** Video of Salactin-HaloTag expressed on top of the native copy visualized by Near-TIRF fluorescent microscopy. Images were taken every 10 seconds for 20 minutes. Video is 150x actual speed. Cyan is Salactin-HaloTag labeled with low concentrations of JF549 to generate speckles, and magenta is Salactin-HaloTag labeled with high concentrations of JF505 to label the whole filament. Scale bar = 2 µm. Movie corresponds to the montage in Figure 2F.

**SM3**. Video of Salactin-HaloTag labeled with JF549 that was expressed on top of the native copy, visualized by Near-TIRF fluorescent microscopy Images were taken every 30 seconds for 30 minutes. Video is 600x actual speed. Scale bar = 4 µm.

**SM4**. Video of Salactin-HaloTag expressed as a sole copy visualized by Near-TIRF fluorescent microscopy. Images were taken every 30 seconds for 1 hour. Video is 600x actual speed. Scale bar = 4 µm.

**SM5**. Video of Salactin-msfGFP expressed as a sole copy visualized by Near-TIRF fluorescent microscopy. Images were taken every 2 minutes for 32 minutes. Video is 1,200x actual speed. Scale bar = 10 µm.

### SUPPLEMENTARY TABLES

**Supplementary Table S1.** Whole genome resequencing data for Δ*salactin*, Δ*ura3*, and NRC-1 wildtype (as a baseline for genomic differences). Each page represents differences from the NRC-1 genome with the specified strain. (see separate excel file)

**Supplementary Table S2.** Distribution of cells with filaments that are dynamic or not

| | WT (*prpa-salactin-msfGFP*) (n = 334) | WT (*prpa-salactin-halotag*) (n=170) | Δ*salactin* (*prpa-salactin-halotag*) (n=129) |
|---|---|---|---|
| Dynamic filaments | 74.55% | 61.18% | 38.76% |
| No dynamics | 25.45% | 38.82% | 61.24% |

**Supplementary Table S3.** The distribution of the different phenotypes in Δ*salactin* (*prpa-salactin-msfGFP*) split between foci, diffuse, and filaments.

| | Δ*salactin* (*prpa-salactin-msfGFP*) (n = 1061) |
|---|---|
| Foci | 93.2% |
| Diffuse | 4.4% |
| Filaments | 2.4% |

**Supplementary Table S4.** Strains used in this study

| Strain | Genotype | Source |
|---|---|---|
| HS148 | Δ*ura3* (*H. salinarum*) | (Darnell et al., 2020) |
| ΔVNG0153C | Δ*salactin* (*H. salinarum*) | Gift from Baliga lab |

| | | |
|---|---|---|
| hsJZ52 | WT (*prpa-salactin-msfGFP*) (*H. salinarum*) | This study |
| hsJZ86 | WT (*prpa-salactin-halotag*) (*H. salinarum*) | This study |
| hsJZ95 | Δ*salactin* (*prpa-salactin-msfGFP*) (*H. salinarum*) | This study |
| hsJZ106 | Δ*salactin* (*prpa-salactin-halotag*) (*H. salinarum*) | This study |

**Supplementary Table S5.** Plasmids used in this study

| Plasmid | Description | Source |
|---|---|---|
| pHS01 | pRPA plasmid | This study (Jenna Eun) |
| pJZHS4 | *prpa-salactin-15aa-msfGFP* | This study |
| pJZHS11 | *prpa-salactin-15aa-halotag* | This study |
| pEJZ2 | *pSUMO-salactin* | This study |
| pEJZ15 | *pSUMO-salactin-GSKCK* | This study |

**Supplementary Table S6.** Primers used in this study

| Primer | Sequence | Description | Source |
|---|---|---|---|
| oJM220 | TCGACGATGTCGATGGTGAC | Used to create qPCR standard curves | This study |
| oJM221 | AGCAAGGATGGGACTGTTCG | Used to create qPCR standard curves | This study |
| oPS48 | CTCGAGCCCGGGTG | Used to amplify linear pSUMO backbone | (Stoddard et al., 2020) |
| oPS49 | ACCACCAATCTGTTCTCTGTG | Used to amplify linear pSUMO backbone | (Stoddard et al., 2020) |
| oHS01 | CGGGTACGCCGAAAGCTTGGATCCGAATTCTGTCGGTTCAGGCCAAGG | Used to amplify rpa promoter (reverse) with a PMTFChis backbone tail (for pHS01) | This study (Jenna Eun) |
| oHS02 | CCCTCCCATGCCACTCTTCACACGCGGTACCTGGCTCCGCAAGCCAAG | Used to amplify rpa promoter (forward) with a PMTFChis backbone tail (for pHS01) | This study (Jenna Eun) |
| oHS263 | GGCCTGAGCCCGGTCCCTGGCCAGATCCCTCGAGGTACTCCCCGAGATCGA | Used to amplify *salactin* (reverse) with a part of the 15 amino acid linker tail (for pJZHS4 and pJZHS11) | This study (Jenna Eun) |
| oJZHs28 | GCCTTGGCCTGAACCGACAGATGTCCGACGATACCGAG | Used to amplify *salactin* (forward) with a rpa promoter tail (for pJZHS4 and pJZHS11) | This study |
| oJZHs44 | GACCGGGCTCAGGCCAAGGTTCCGGCCGAAAAGGGGAAGAATTG | Used to amplify msfGFP (forward) with a part of the 15 amino acid linker tail (for pJZHS4) | This study |
| oJZHs48 | CCGAAAGCTTGGATCCGCTATCATTTGTAAAGTTCATCCATTC | Used to amplify msfGFP (reverse) with a prpa backbone tail (for pJZHS4) | This study |

| | | | |
|---|---|---|---|
| oJZHs57 | CAGAGAACAGATTGGTGGTATGTCCGACGATACCG | Used to amplify *salactin* (forward) with a pSUMO backbone tail (for pEJZ2 and pEJZ15) | This study |
| oJZHs58 | CCCGGGCTCGAGCTAGTACTCCCCGAGATC | Used to amplify *salactin* (reverse) with a pSUMO backbone tail (for pEJZ2) | This study |
| oJZHs87 | CGGGCTCAGGCCAAGGTTCCGGCGCAGAAATCGGTACTGGC | Used to amplify HaloTag (forward) with a part of the 15 amino acid linker tail (for pJZHS11) | This study |
| oJZHs88 | GAAAGCTTGGATCCGCTATTAGCCGCTGATTTCTAAGGT | Used to amplify Halotag (reverse) with a prpa backbone tail (for pJZHS11) | This study |
| oJZHs118 | CACCCGGGCTCGAGCTATTTGCATTTGCTGCCGTACTCCCCGAGATCG | Used to amplify *salactin* (reverse) with a GSKCK+pSUMO backbone tail (for pEJZ15) | This study |

### SUPPLEMENTARY METHODS – PLASMID CONSTRUCTION

**pHS01**: Modified pMTFChis (Darnell et al., 2020), with a rpa (gene locus tag: VNG0133G (old), VNG_RS00545) promoter instead of a fdx (gene locus tag: VNG2293G (old), VNG_RS00545) promoter was generated with two fragments: 1) the RPA promoter (with pMTFChis backbone overhang) was PCR amplified using oHS01 and oHS02 from the NRC-1 *H. salinarum* genome, 2) the plasmid backbone created by cutting the pMTFChis plasmid with KpnI and EcoRI restriction enzmyes. The two pieces were assembled using Gibson assembly (Gibson, 2011).

**pJZHS4**: *prpa-salactin-15aa-msfGFP* was generated with three fragments: 1) *salactin* (with plasmid backbone and 15 amino acid linker (15aa) overhang), which was PCR amplified using oHS263 and oJZHs28 from the NRC-1 genome, 2) msfGFP (with plasmid backbone and 15aa overhang), which was PCR amplified using oJZHs44 and oJZHs48 from a gBlock gene fragment obtained from Dion and colleagues (Dion et al., 2019), 3) linear pHS01 backbone, which was made with restriction enzyme digest with EcoRI. The three pieces were assembled using Gibson assembly (Gibson, 2011).

**pJZHS11**: *prpa-salactin-15aa-halotag* was generated with three fragments: 1) *salactin*, which was PCR amplified using oHS263 and oJZHs28 from the NRC-1 genome or pJZHS4, 2) HaloTag, which was PCR amplified using oJZHs87 and oJZHs88 from a gBlock gene fragment obtained from Dion and colleagues (Dion et al., 2019), 3) linear pHS01 backbone, which was made with restriction enzyme digest with EcoRI. The three pieces were assembled using Gibson assembly (Gibson, 2011).

**pEJZ2**: *pSUMO-salactin* (his6-SUMO tagged Salactin in a T7 expression vector) was generated with two fragments: 1) *salactin* (with pSUMO backbone overhang), which was PCR amplified using oJZHs57 and oJZHs58 from the NRC-1 genome or pJZHS4, 2) linear pSUMO backbone, which was PCR amplified using oPS48 and oPS49. The two pieces were assembled using Gibson assembly (Gibson, 2011).

**pEJZ15**: *pSUMO-salactin-GSKCK* was generated with two fragments: 1) *salactin* (with pSUMO backbone overhang and addition of the GSKCK), which was PCR amplified using oJZHs57 and oJZHs118 from the NRC-1 genome or pJZHS4, 2) linear pSUMO backbone, which was PCR amplified using oPS48 and oPS49. The two pieces were assembled using Gibson assembly (Gibson, 2011).
